# A truncating mutation of *Magel2* in the rat modelled for the study of Schaaf-Yang and Prader-Willi syndromes alters select behavioral and physiological outcomes

**DOI:** 10.1101/2022.08.09.503377

**Authors:** Derek L Reznik, Mingxiao V Yang, Pedro Albelda de la Haza, Antrix Jain, Melanie Spanjaard, Susanne Theiss, Christian P Schaaf, Anna Malovannaya, Theresa V Strong, Surabi Veeraragavan, Rodney C Samaco

## Abstract

Truncating mutations of the maternally imprinted, paternally expressed *MAGEL2* gene are the predicted genetic cause of several rare neurodevelopmental disorders including Schaaf-Yang (SYS), Chitayat-Hall and Opitz Trigonocephaly C syndromes. *MAGEL2* is also deleted or inactivated in Prader-Willi syndrome (PWS). Previous studies in mice have utilized *Magel2* gene deletion models to examine the consequences of its absence. In this study, we report the generation, molecular validation, and phenotypic characterization of a novel rat model with a truncating *Magel2* mutation generating a mutant peptide sequence more closely modeling variants associated with SYS-causing mutations. Within the hypothalamus, a brain region wherein mouse and human MAGEL2 is paternally-expressed, we demonstrate at the level of transcript and peptide detection that *Magel2* in the rat exhibits a paternal, parent-of-origin effect. In the evaluation of behavioral features across several domains, juvenile *Magel2* mutant rats display select alterations in anxiety-like behavior and sociability measures. Moreover, the analysis of peripheral organ systems detected alterations in body composition, cardiac structure and function, and breathing irregularities in *Magel2* mutant rats. Several of these findings are concordant with reported mouse phenotypes, signifying the conservation of MAGEL2 function across rodent species for specific behavioral outcome measures. We conclude that our comprehensive analysis demonstrating impairments across multiple domains demonstrates the tractability of this model system for the study of truncating *MAGEL2* mutations.

## INTRODUCTION

The neurodevelopmental disorders (NDD) Schaaf-Yang (SYS, OMIM # 615447), Chitayat-Hall (CHS) and Opitz Trigonocephaly C syndromes (OTCS, OMIM 211750) share overlapping clinical features that have been attributed to commonly shared loss-of-function truncating mutations in the imprinted gene *MAGEL2*. MAGEL2 is enriched in the brain, heterozygous mutations in the active paternal copy of *MAGEL2* leave individuals vulnerable to neurological dysfunction as the maternal allele is epigenetically silenced (Boccaccio et al., 1999; Schaaf et al., 2013). Previous studies into the consequences of MAGEL2 deficiency have focused on its complete loss-of-function, as it is also a gene frequently deleted in patients with Prader-Willi syndrome (PWS, OMIM #176270).

Mice carrying deletions of the paternal *Magel2* have been the primary mammalian system used to examine the biological systems and molecular pathways effected in the absence of MAGEL2 (Bervini and Herzog, 2013). While the current murine deletion models may be effective tools to study the consequences of the complete absence of MAGEL2, they may also limit our knowledge in the evaluation of consequences that are directly linked to *MAGEL2* truncating mutations. Moreover, due to an increased prevalence of intellectual disability, autism spectrum disorder (ASD), and disrupted social behaviors in people with MAGEL2-related disorders including SYS and PWS (McCarthy et al., 2018; Patak et al., 2019; Marbach et al., 2020), evaluating the consequences of truncated MAGEL2 in species that enable the study of more complex behavioral patterns may provide deeper insight for the study of these specific phenotypes.

One approach to complement studies of mouse models for understanding rare NDD is the laboratory rat, a genetically tractable system with greater genetic conservation to the human genome (Gibbs et al., 2004). The behavioral repertoire of the rat includes sophisticated social and cognitive behaviors (Pellis and Pellis, 1998; Rudy, 2009). The innate difference in sociability compared to mice (Pellis and Pasztor, 1999) presents a unique opportunity for research into the behavioral features of NDD. Beyond direct consequences of the central nervous system (CNS), people with NDD also frequently suffer from dysfunction of peripheral organ systems. In the added context of therapeutic development, the large overall size of the rat offers unique advantages in preclinical proof-of-concept validation studies (Wöhr and Scattoni, 2013); combined with techniques that are readily accessible to manipulate the rat genome, the laboratory rat enables a level of study that can enhance ongoing efforts for the study of and development of treatments for rare genetic NDD. To this end, the purpose of our study was to identify the neurobehavioral, organ system and molecular consequences of a truncating *Magel2* mutation in the rat. By using an evolutionarily more complex mammalian system than the mouse, we sought out to identify changes in surrogate phenotypic domains that may be pathogenically relevant to mechanistic pathways underlying the consequences of *MAGEL2* truncation mutations.

## RESULTS

### Transcription activator-like effector nuclease (TALEN) targeting of rat *Magel2* results in a predicted truncating mutation that does not impact levels of mRNA abundance

An eight-base pair deletion was generated in the single exon coding sequence of the rat *Magel2*, c.735_742del, and was confirmed with sequencing. The deletion results in a frameshift event causing a change from serine to glycine at amino acid position 132, in turn a mutant peptide sequence extends through to a premature termination site at amino acid 728 resulting in a predicted truncated MAGEL2 (**Fig. 1A**). Given that *Magel2* is enriched in the hypothalamus, we focused on its expression within this brain region. To enable a complete readout of total mRNA abundance, we used primers that spanned a region after the TALEN-targeted mutation site in *Magel2* to amplify a product commonly shared between the mutant and wildtype alleles independent of potential MAGEL2 parent-of-origin effects. Relative mRNA abundance was comparable in the hypothalami of rats representing all four possible *Magel2* genotypes. Heterozygous rats with either the paternally- or maternally-inherited *Magel2* TALEN-generated mutation (*Magel2^m+/p-^*, indicated as m+/p-; and *Magel2^m-/p+^*, indicated as m-/p+) as well as homozygous rats with bi-parental inheritance of the *Magel2* TALEN-generated mutation (*Magel2^m-/p-^*) showed normal *Magel2* mRNA abundance in comparison to wildtype rats (**Fig. 1B**).

**Fig. 1.**
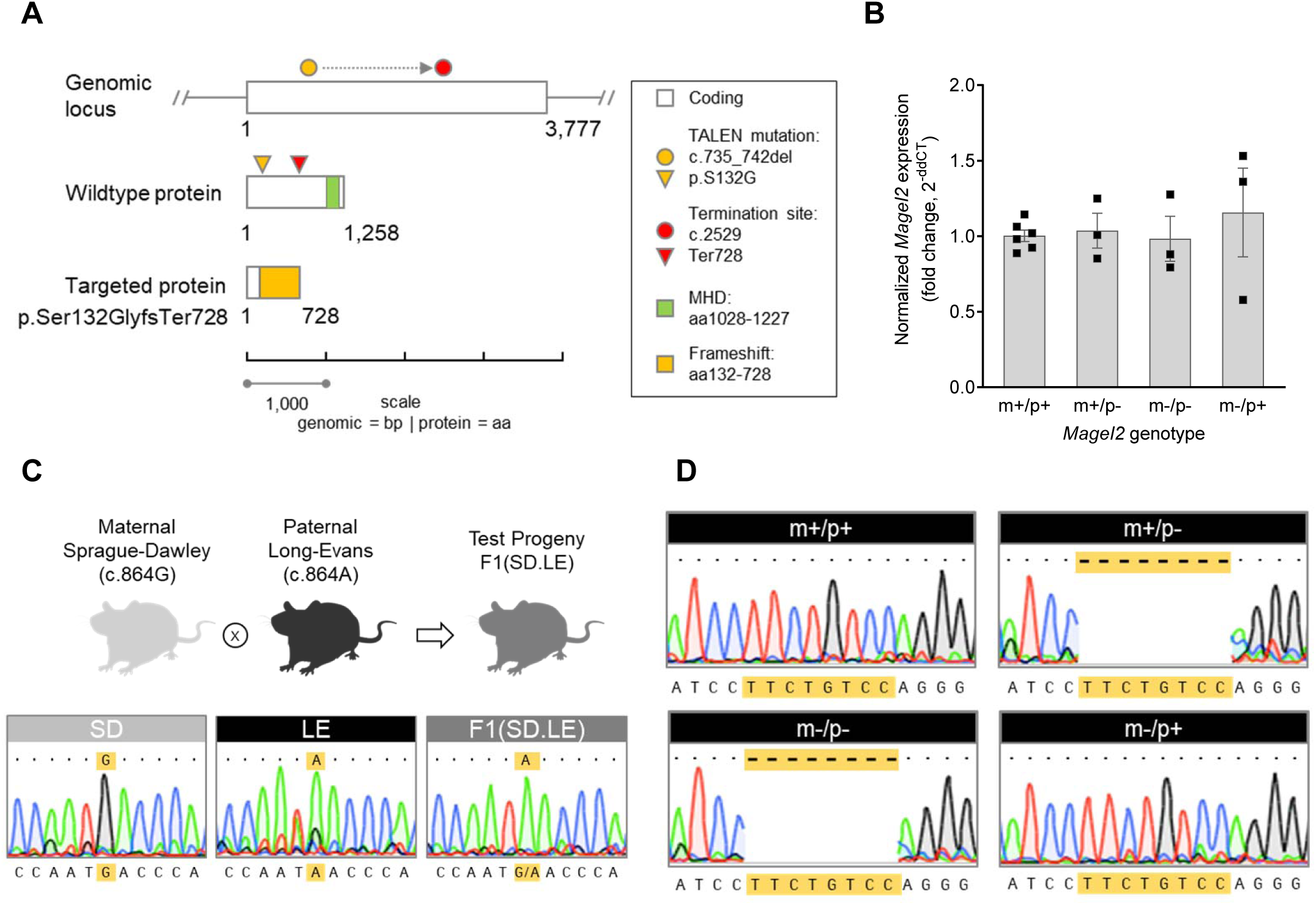
Strategy and validation of a novel *Magel2* rat model. **(A)** Schematic diagram showing the overall strategy to create a *Magel2* rat model. Rat *Magel2* is a single exon gene located on chromosome 1 (genomic locus, white bar; 3,777 bp), encoding a protein product (wildtype protein, grey bar; 1,258 aa) with a MAGE Homology Domain (MHD, green bar). Transcription activator-like effector nuclease (TALEN) modification of the wildtype genomic locus created an eight base pair deletion (c.735_742; gold circle), predicted to shift the open reading frame and result in loss of full-length MAGEL2 protein (targeted protein, p.Ser132Glyfs728). Additional predicted features of the 8bp-induced mutation include translational of a novel amino acid sequence (targeted protein, gold bar) and loss of the conserved MHD owing to introduction of a premature stop codon at position aa728 (targeted protein, red bar). **(B)** Relative *Magel2* mRNA abundance measured by qRT-PCR showed no difference in *Magel2* expression levels in mutant genotype groups compared to wildtype littermate rats. Allelic inheritance and corresponding genotype of each *Magel2* group is shown as either maternal (m) or paternal (p), and wildtype (+) or mutant (-). Comparative analyses were conducted using hypothalamic tissue samples from rats heterozygous for the paternally-inherited or maternally-inherited mutant *Magel2* allele (m+/p- or m-/p+; respectively), rats homozygous for both the paternally-inherited and maternally-inherited mutant *Magel2* alleles (m-/p-), and wildtype animals (m+/p+). Data depicted as mean +/- SEM; individual data points are denoted as black squares N=3-6/rats per genotype group. **(C)** Endpoint RT-PCR and sequencing of hypothalamic cDNA from parental rat strains (maternal Sprague-Dawley, SD; paternal Long-Evans, LE) and F1 hybrid progeny (F1.SD.LE) confirms a parent-of-origin effect on rat *Magel2* expression. The wildtype *Magel2* coding sequence contains a SNP at nt position c.864 that differs between Sprague-Dawley (SD, c.864G) and Long-Evans (LE, c.864A). Representative sequence traces of the SNP from parental and F1 hybrid genotypes is shown (gold highlight); F1(SD.LE) hybrid rats generated from an intercross of maternal SD rats and paternal LE rats show expression of paternal LE-derived SNP, c.864A. Rat images created with BioRender.com. **(D)** Endpoint RT- PCR and sequencing of hypothalamic cDNA from rats derived from the *Magel2* mutant model validates a parent-of-origin effect on rat *Magel2* expression in the context of the novel TALEN-modified *Magel2* rat allele. Comparative analyses were conducted using hypothalamic tissue samples from all four *Magel2* rat genotype groups (wildtype littermates, m+/p+; uniparentally-inherited heterozygous *Magel2* rats, m+/p- and m-/p+; and biparentally-inherited *Magel2* homozygous rats, m-/p-). Representative sequence traces show each *Magel2* genotype group test sequence aligned to the rat reference sequence. Matching sequences (dots) in each window are denoted. Wild-type (m+/p+) and maternal heterozygous *Magel2* rats (m-/p+) show complete alignment to the reference sequence. The absence of sequence (dash, highlighted in gold) corresponding to the TALEN-generated eight bp deletion was found only in *Magel2* genotype groups with paternal inheritance of the *Magel2* mutation (m+/p- and m-/p- rats).

### Rat *Magel2* transcript undergoes parent-of-origin specific expression

Several rat loci are known to be imprinted (Berg et al., 2020; Overall et al., 1997); however, rat *Magel2* has not been evaluated in this species even though it is maternally imprinted in both mice and humans (Boccaccio et al., 1999). Given the absence of altered mRNA abundance in the *Magel2* TALEN-generated rat model (**Fig. 1**), we reasoned that an evaluation of parent-of-origin effects would yield important control evidence to test the validity of the rat model. Therefore, to determine if *Magel2* is paternally-expressed in the rat, we leveraged a naturally occurring single nucleotide polymorphism (SNP) identified between two strains of rats at nucleotide (nt) position 864 (Rat Genome Database, Rnor 5.0, Smith et al. 2020) to enable an analysis of parent-of-origin expression in progeny derived from Long-Evans (LE) male rats mated to Sprague-Dawley (SD) female rats. The 864^th^ nucleotide in SD rats is a guanine, while LE rats display an adenine. By amplifying a product spanning this region followed by Sanger sequencing, we found that hypothalamic *Magel2* shows a pattern of paternal inheritance and expression in F1 hybrid test progeny, indicated by an adenine at nt position 864 (**Fig. 1C**). Using the identical endpoint RT-PCR approach, we tested whether the *Magel2* mutant rats in either the heterozygous or homozygous mutation state showed a similar parent-of-origin, paternal-specific expression pattern. cDNA sequencing of mutant and wildtype hypothalami showed that the *Magel2* mutation is indeed preferentially expressed from the paternal allele, as evident in *Magel2^m+/p-^* and *Magel2^m-/p-^* progeny with paternal inheritance of the *Magel2* mutation (denoted as the genotypes m+/p- and m-/p-; respectively, **Fig. 1D**). In comparison, chromatograms from *Magel2^m+/p+^* progeny with inheritance of inheritance of wildtype parental *Magel2* alleles, and *Magel2^m-/p+^* with maternal inheritance of the *Magel2* mutation (denoted as the genotypes m+/p+ and m-/p+; respectively, **Fig. 1D**), showed normal cDNA sequence traces. Taken together, these controls studies confirm that *Magel2* is preferentially expressed from the paternally-inherited allele across all genotypes. Moreover, expression from the maternally-inherited allele in the rat hypothalamus was undetectable by Sanger sequencing, suggesting that the total mRNA abundance measured by QRT-PCR reflected expression levels specifically from the paternally-expressed *Magel2*.

### Paternal heterozygous or homozygous loss of *Magel2* in juvenile rats results in altered perseverative-repetitive behavior, anxiety-like behavior, and sociability

The primary goal of our phenotype profiling aimed to test the hypothesis that rats with the paternally-inherited *Magel2* mutation may be a construct valid model to study outcomes that emerge as a consequence of truncated MAGEL2. To this end, independent cohorts were generated to first evaluate test progeny derived from breeding schemes producing rats with the paternally-inherited *Magel2* mutation (*Magel2^m+/p-^*, denoted as m+/p-), followed by examination of homozygous, bi-parental inheritance of the *Magel2* mutation (*Magel2^m-/p-^*, denoted as m-/p-). Wildtype littermates were also generated within each breeding scheme and evaluated as control comparisons. To test the neurobehavioral consequences of the truncating MAGEL2 mutations in the rat, we deployed a well-defined battery of test procedures to assess for alterations across multiple domains including anxiety-like behavior, sociability, locomotor function, sensorimotor gating, perseverative behavior, learning and memory and pain nociception (Veeraragavan et al., 2016).

*Magel2^m+/p-^* rats displayed significantly altered behavioral outcomes in select social and perseverative behaviors (**Fig. 2**). In the elevated circle maze test for anxiety-like behavior (Braun et al., 2011), *Magel2^m+/p-^* rats in comparison to wildtype littermates showed normal exploratory activity in the open and enclosed regions elevated circle maze (**Fig. 2A**). In the three chamber test for social approach (Ku et al., 2016; Veeraragavan et al., 2016; Yang et al., 2011), the typical expected pattern of social approach behavior was observed by spending significantly more time investigating a novel conspecific partner compared to a novel object (**Fig. 2B**). To further investigate sociability, we evaluated dyadic direct social play behaviors. In contrast to the laboratory mouse which has been reported to display only rudimentary forms of play-like behavior, direct social interaction with dyad pairs of rats enables the study of developmentally-restricted social behaviors uniquely observed in juvenile aged rats (Pellis and Pellis, 1998; Veeraragavan et al., 2016). We quantified play-like behavior, sniffing and following behavior, and general contact defined as test subject paw to conspecific subject torso (paw-to-torso contact) interaction in dyad interactions between either mutant or wildtype rats, with wildtype non-littermate same-sex rat conspecific partners. Among these three behavioral categories, we found that *Magel2^m+/p-^* rats in comparison to wildtype littermates showed no difference in the duration of play-like behavior, a significant increase in the duration of sniffing and following conspecific wildtype non-littermate partner rats, and normal duration of general contact behaviors (**Fig. 2C-E**). Similarly, the number of observed events during juvenile dyadic interactions was comparable across genotypes for play-like and general contact behaviors, and significantly increased for sniffing and following behavior in *Magel2^m+/p-^* rats in comparison to wildtype littermates (**Supplementary Material, Figure S1A-C**). To evaluate perseverative, repetitive-like behavior, we examined the performance of *Magel2^m+/p-^* rats and their wildtype littermates in the marble burying test and in an overnight assessment of the block chew test (Hamilton et al., 2014; Veeraragavan et al., 2016). *Magel2^m+/p-^* rats displayed a significant reduction in the number of marbles buried (**Fig. 2F**), with no difference in the block chew test (**Fig. 2G**). To measure spontaneous locomotor exploratory activity in a novel environment, *Magel2^m+/p-^* rats in comparison to wildtype littermates showed a modest but significant decrease in total distance traveled in the open field assay (**Fig. 2H**). Other measures of generalized locomotor activity in the open field test including center distance traveled, vertical activity, and the ratio of center-to-total distance traveled were comparable across genotype groups (**Supplementary Material, Fig. S1D-F**). To evaluate sensorimotor gating, we tested prepulse inhibition of the acoustic startle response in rats presented with a randomized series of low decibel sound levels immediately preceding a loud decibel sound (120 dB). *Magel2^m+/p-^* rats displayed a significant reduction in baseline startle reactivity to the intense decibel sound (**Fig. 2I**); however, prepulse inhibition of the startle response was normal (**Fig. 2J**).

**Fig. 2.**
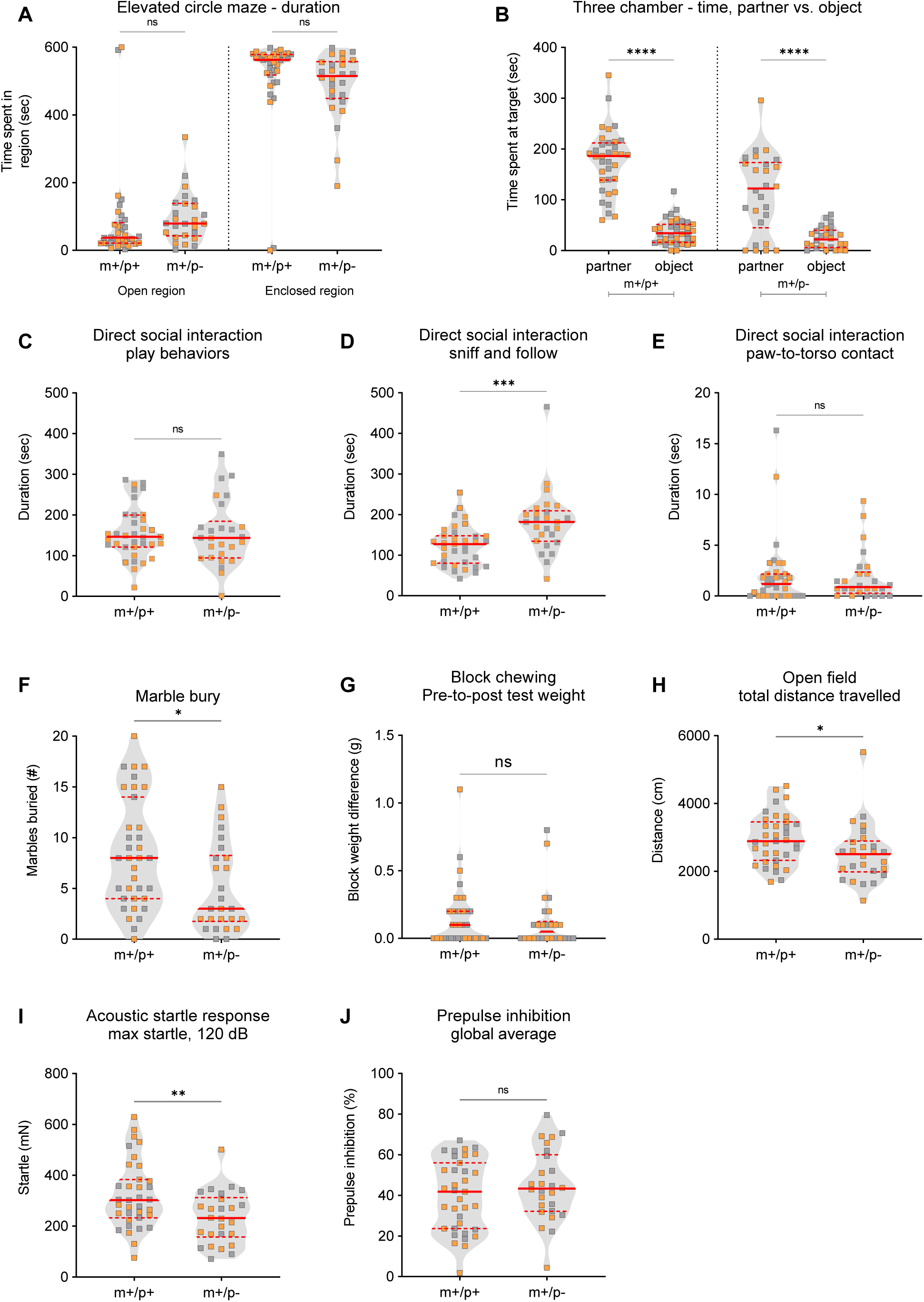
*Magel2^m+/p-^*rats display select alterations in dyadic social interaction, perseverative-like behavior, locomotor activity and startle response. (**A**) *Magel2^m+/p-^* rats (m+/p-) compared to WT littermates (m+/p+) spent a comparable amount of time in both the open and enclosed regions of the elevated circle maze. (**B**) *Magel2^m+/p-^* rats and WT littermates spent a statistically significant amount of time investigating conspecific partners in comparison to the novel object in the three-chamber test for social approach. **(C-E)** In a test of juvenile dyad social interaction, *Magel2^m+/p-^* rats in comparison to WT littermates show a comparable amount of time engaged in play-like behavior **(C)**, a statistically significant increase in the duration of sniffing and following conspecific partners **(D)**, and a comparable time spent in general paw-to-torso contact **(E)**. **(F, G)** In tests of perseverative-like behavior, *Magel2^m+/p-^* rats bury fewer marbles in comparison to WT littermates **(F)**; no differences were observed in the block chew test **(G)**. (H) *Magel2^m+/p-^* rats showed a significant decrease in total distance traveled. **(I, J)** Acoustic startle response at 120dB in *Magel2^m+/p-^* rats was significantly reduced **(I)**; prepulse inhibition of the startle response was normal **(J)**. Violin plots were used to indicate data density and distribution, shown are grey squares representing males and orange squares representing females within each genotype/group category. Solid red line and dashed red lines indicate median and quartiles; respectively. Asterisks indicate statistical significance of mutant rats relative to wildtype littermate controls. *p<0.05 **p<0.01; ns, not significant. Statistical summary of behavioral data are provided in **Supplementary Material, Table ST1**.

Homozygous *Magel2^m-/p-^* rats were also tested in the identical test battery to determine if bi- parental inheritance of the truncating mutation alters behavioral outcomes. We found that *Magel2^m-/p-^* rats exhibited altered anxiety-like behaviors and sociability. In the elevated circle maze, *Magel2^m-/p-^* rats spent more time in the open regions (**Fig. 3A**) less time in the closed regions (**Fig. 3A**) and transitioned more frequently between the two areas (**Supplementary Material, Fig. S2A**). In the three chamber test for indirect social interaction, a repeated measures two-ANOVA indicated a significant genotype-by-sex interaction in three test parameters (**Supplementary Material, Table ST1**). *Post hoc* comparisons revealed that male *Magel2^m-/p-^* rats showed a comparable amount of time spent investigating a novel partner versus novel object, suggesting altered social approach behavior (**Fig. 3B**). Similar findings were observed in the number of events, or visits, to the partner versus object, as well as in the number of events to the chamber sides containing either the partner or object (**Supplementary Material, Fig. S2B, C**). These parameters were not altered in male or female wildtype littermates or female *Magel2^m-/p-^* rats, suggesting a male-specific social approach phenotype in rats homozygous mutant for the truncating *Magel2* mutation. To deepen our analysis of sociability, we deployed the juvenile dyadic direct social interaction test. Significant genotype-by-sex interactions followed by post hoc analyses revealed a similar pattern of deficits in *Magel2^m-/p-^* rats. The time spent and number of events engaged in play-like behavior was significantly reduced in male *Magel2^m-/p-^* rats by time spent playing (**Fig. 3C, D**). Sniff-follow duration was normal (**Fig. 3E**); however, unlike other parameters, the number of events observed was significantly reduced in both male and female *Magel2^m-/p-^* rats (**Fig. 3F**). In contrast to rats heterozygous for the paternally-inherited *Magel2* mutation, *Magel2^m-/p-^* rats did not display altered perseverative behaviors in either the marble burying or block chewing tests (**Fig. 3G, H**). Performance in the open field test was normal (**Fig. 3I, Supplementary Material, Fig. S2**); therefore the decreased anxiety-like behavior in the elevated circle maze and altered social behavior does not appear to be a result of alterations in gross locomotor function. Finally, in the evaluation of prepulse inhibition of the acoustic startle response, we found *Magel2^m-/p-^* rats displayed normal baseline startle reactivity and normal PPI (**Fig. 3J, K**).

**Fig. 3.**
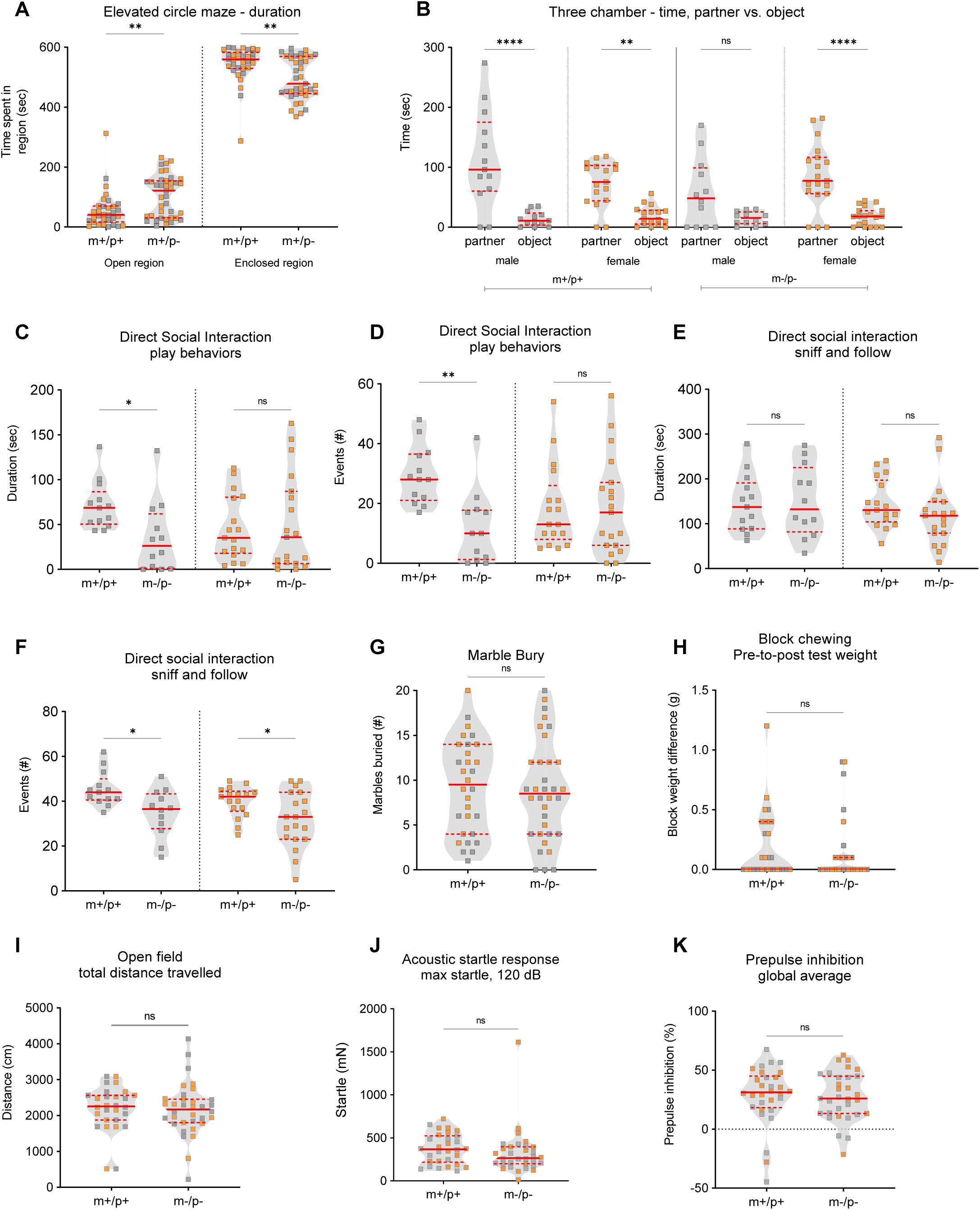
*Magel2^m-/p-^*rats display select alterations in anxiety-like behavior, social approach and dyadic social interaction. **(A)** *Magel2^m-/p-^* (m-/p-) compared to WT littermates (m+/p+) spent more time in the open regions, less time in the closed regions in the elevated circle maze. **(B)** Wildtype male and female littermates, and *Magel2^m-/p-^* female rats spent more time investigating a partner versus object in the three-chamber test as expected for typical developing animals; in contrast, the time *Magel2^m-/p-^* rats investigated the partner versus object was not significant. **(C-F)** In dyadic social interaction tests, male but not female *Magel2^m-/p-^* rats showed a reduction in play-like behavior duration and number of events (C, D). In addition, while male and female *Magel2^m-/p-^* rats showed normal sniff and follow duration **(E)**, both male and female *Magel2^m-/p-^* rats showed a significantly reduced number of sniff and follow events **(F) (G, H)** In tests for perseverative-like behavior, *Magel2^m-/p-^* rats showed no significant differences in marble burying **(G)** or block chewing **(H)**. **(I)** Total distance traveled in the open field was comparable across genotypes. **(J, K)** Acoustic startle reactivity to a loud decibel noise (120dB) **(J)** and prepulse inhibition of the startle response **(K)** was normal in *Magel2^m+/p-^* rats. Violin plots were used to indicate data density and distribution, shown are grey squares representing males and orange squares representing females within each genotype/group category. Solid red line and dashed red lines indicate median and quartiles; respectively. Asterisks indicate statistical significance of mutant rats relative to wildtype littermate controls. *, *p* <0.05; ** *p* < 0.01; ns, not significant. Statistical summary of behavioral data are provided in **Supplementary Material, Table ST1**.

### Paternal heterozygous or homozygous loss of *Magel2* in juvenile rats does not affect memory and learning performance or pain nociception

PWS and SYS patients are frequently diagnosed with intellectual disability and developmental delay. To determine if mutant rats for the truncating *Magel2* mutation exhibited deficits in learning and memory, *Magel2^m+/p-^* and *Magel2^m-/p-^* rats were tested in object recognition memory and aversive fear memory. Neither *Magel2^m+/p-^* nor *Magel2^m-/p-^* rats displayed observable deficits in the novel object recognition index (**Supplementary Material, Fig. S3A, B**) or in contextual or cue-based memory retrieval (**Supplementary Material, Fig. S3C-F**. As a relevant control measure for the mild shock-based conditioned fear testing paradigm and to test whether alterations in pain sensitivity may be present in *Magel2* mutant rats, thermal pain reactivity to a 55°C heated surface was measured. No genotype differences were observed genotypes in the latency to first response in the hot plate test (**Supplementary Material, Fig. S3G, H**). A heatmap (**Fig. 4**) summarizing the degree of statistical significance for behavioral outcomes identified in each *Magel2* mutant test cohort relative to their respective wildtype littermates is provided, analyzed by repeated measures two-way ANOVA (**Fig. 4A**) or multivariate two-way ANOVA (**Fig. 4B**). Corresponding statistical summary data are provided in **Supplementary Material, Table ST1.**

**Fig. 4.**
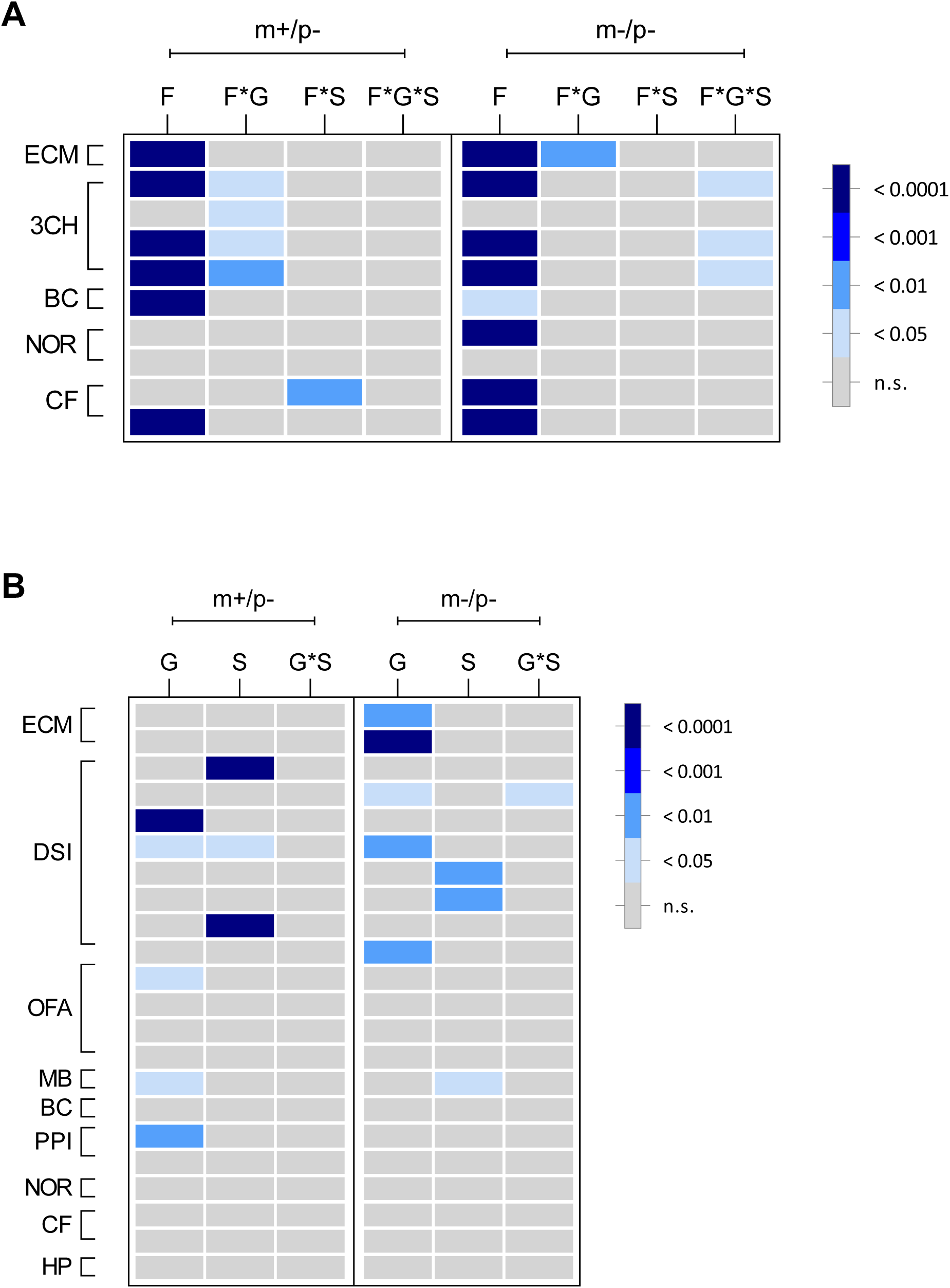
Summary heatmap representation of mutant *Magel2* rat behavioral outcomes. **(A, B)** Graphical representation showing an overview of the degree of statistical significance for both mutant *Magel2* lines studied relative to their wildtype littermate controls. Each row corresponds to a specific test parameter of the indicated procedure to the left outlined in brackets. Within each mutant line studies (shown along top of graph as m+/p- or m-/p-) each column represents the main effect or interaction for each statistical analysis. As shown in each panel legend, the degree of statistical significance is indicated by relative grey to dark blue color, with each color corresponding to a P value ranging from not significant (ns) to p < 0.0001. **(A)** For applicable procedures, repeated measures ANOVA was conducted for specific parameters of the elevated circle maze (ECM), three chamber test (3CH), block chew test (BC), novel object recognition test (NOR) and conditioned fear (CF). The specific parameters for each row are also indicated in **Supplementary Material, Table ST1**; F = factor, F*G = factor-by-genotype, F*S =factors-by-sex, F*G*S = factor-by-genotype-by-sex. **(B)** For parameters that were not analyzed by RM-ANOVA, multi-variate ANOVA using genotype and sex as main effects followed by post-hoc analyses was conducted; p values were graphically displayed using the same heatmap notation as described above. Elevated circle maze (ECM), three chamber test (3CH), direct dyadic social interaction (DSI), open field assay (OFA), marble bury test (MB), block chew test (BC), acoustic startle and prepulse inhibition (PPI), novel object recognition test (NOR) and conditioned fear (CF), hot plate (HP). The specific parameters for each row are also indicated in **Supplementary Material, Table ST1**; G = genotype, S = sex, G*S = genotype-by-sex.

### Body composition and weight are altered in paternal and homozygous *Magel2* mutant rats

Individuals with *MAGEL2* variants exhibit several structural and functional alterations of peripheral organ systems; these features have been reported to co-occur with behavioral phenotypes (Patak et al., 2019). Therefore, to evaluate whether disruption of rat *Magel2* manifests physiological alterations that co-occur with behavioral phenotypes, we set out to test additional measurements of cardiac, skeletal, and respiratory systems. Body morphological content such as muscle and fat percentages were performed to examine the tissue content and overall weight differences. Both *Magel2^m+/p-^* and *Magel2^m-/p-^* rats exhibited decreased total body weight at PND21 in comparison to their wildtype littermates (**Fig. 5A, B**). At 4 weeks of age, *Magel2^m+/p-^* rats continued to display decreased total weight, while *Magel2^m-/p-^* rats appeared to show no difference at the same age (**Fig. 5C, D**). Weight differences continued to persist at 12 weeks of age; however at 36 weeks of age a significant genotype-by-sex interaction followed by post-hoc analyses showed that only male *Magel2^m+/p-^* rats showed a weight difference compared to their respective male wildtype littermates (**Fig. 5E, F**). Evaluation of lean and fat mass at 4 weeks of age identified a decrease in lean but not fat mass at 4 weeks of age in *Magel2^m+/p-^* rats; a similar analysis of lean and fat mass at 12 weeks of age did not reveal any significant group differences (**Fig. 5G, H**). In addition to altered total body weight and lean weight, *Magel2^m+/p-^* rats displayed significantly decreased bone area, mineral content and density at 4 weeks of age (**Fig. 6A-C**).

**Fig. 5.**
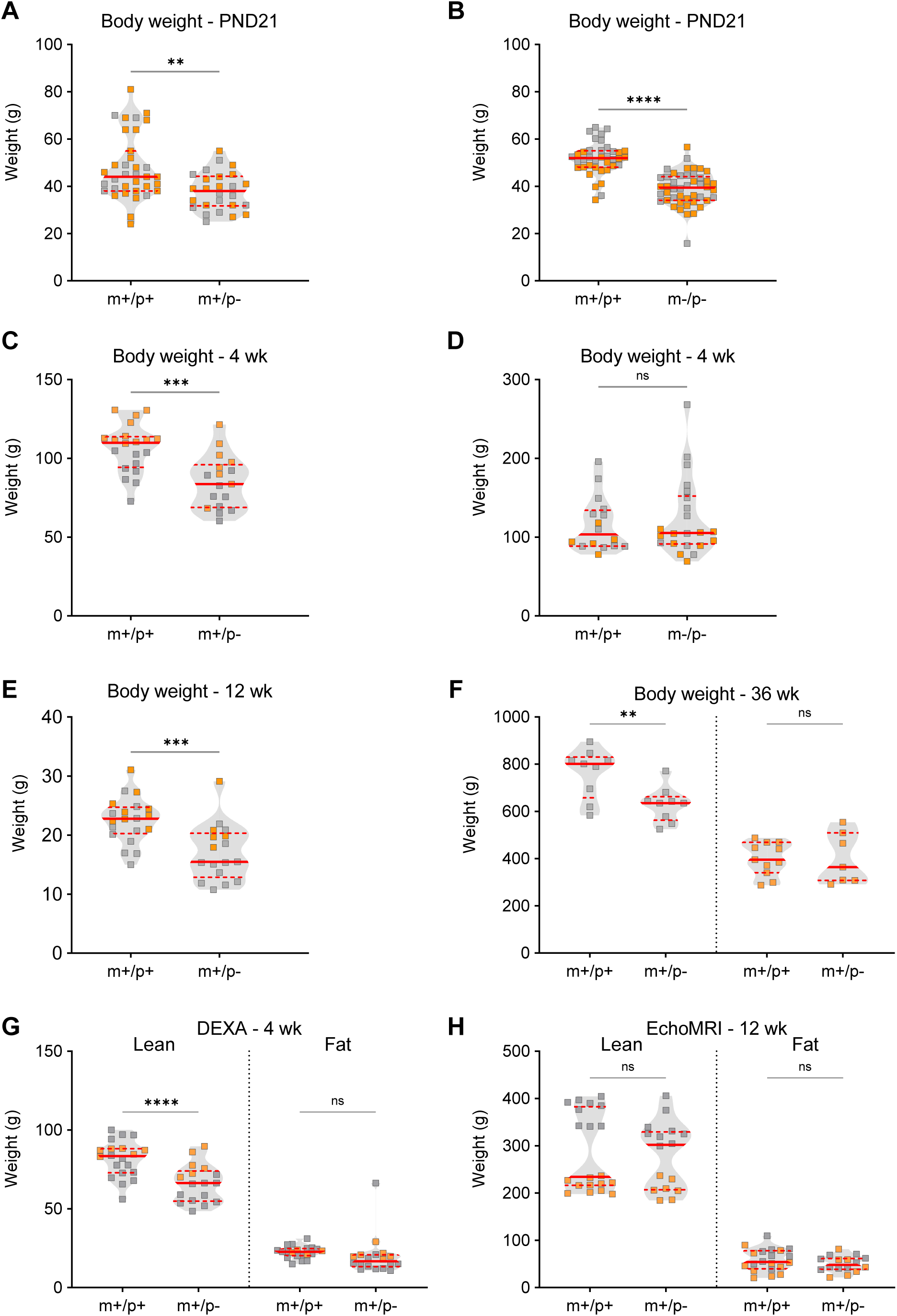
Alterations in body weight and composition were found in rats with paternally-inherited *Magel2* mutation. **(A-D)** For *Magel2^m+/p-^* (m+/p-) rats and their respective wildtype littermates (m+/p+), and for *Magel2^m-/p-^* rats (m-/p-) rats and their respective wildtype littermates (m+/p+) body weight was assessed at PND21, and at 4 weeks of age. At PND21, both m+/p- and m/p- rats showed a significant reduction in weight compared to their respective m+/p+ littermates **(A, B)**; in contrast, at 4 weeks of age, m+/p- rats but not m-/p- rats showed a significant reduction in body weight **(C, D)**. **(E, F)** *Magel2^m+/p-^* (m+/p-) rats showed decreased body weight at 12 weeks of age **(E);** at 36 weeks of age only male m+/p- rats showed a reduction in body weight **(F)**. **(G, H)** Body composition for measurements of lean and fat mass was evaluated by DEXA at 4 weeks of age and MRI at 12 weeks of age. At 4 weeks of age *Magel2^m+/p-^* (m+/p-) rats showed a significant reduction in lean but not fat mass **(G)**; at 12 weeks of age, the absence of a genotype-by-sex interaction resulted in an overall genotype group analysis; no differences were observed between m+/p- and their respective m+/p+ littermates **(H)**. Violin plots were used to indicate data density and distribution, shown are grey squares representing males and orange squares representing females within each genotype/group category. Solid red line and dashed red lines indicate median and quartiles; respectively. Asterisks indicate statistical significance of mutant rats relative to wildtype littermate controls. * *p* < 0.05; ** *p* < 0.01; *** *p* < .001; **** *p* < 0.0001; ns, not significant; two-way ANOVA with genotype and sex as factors, followed by post-hoc analyses as appropriate.

**Fig. 6.**
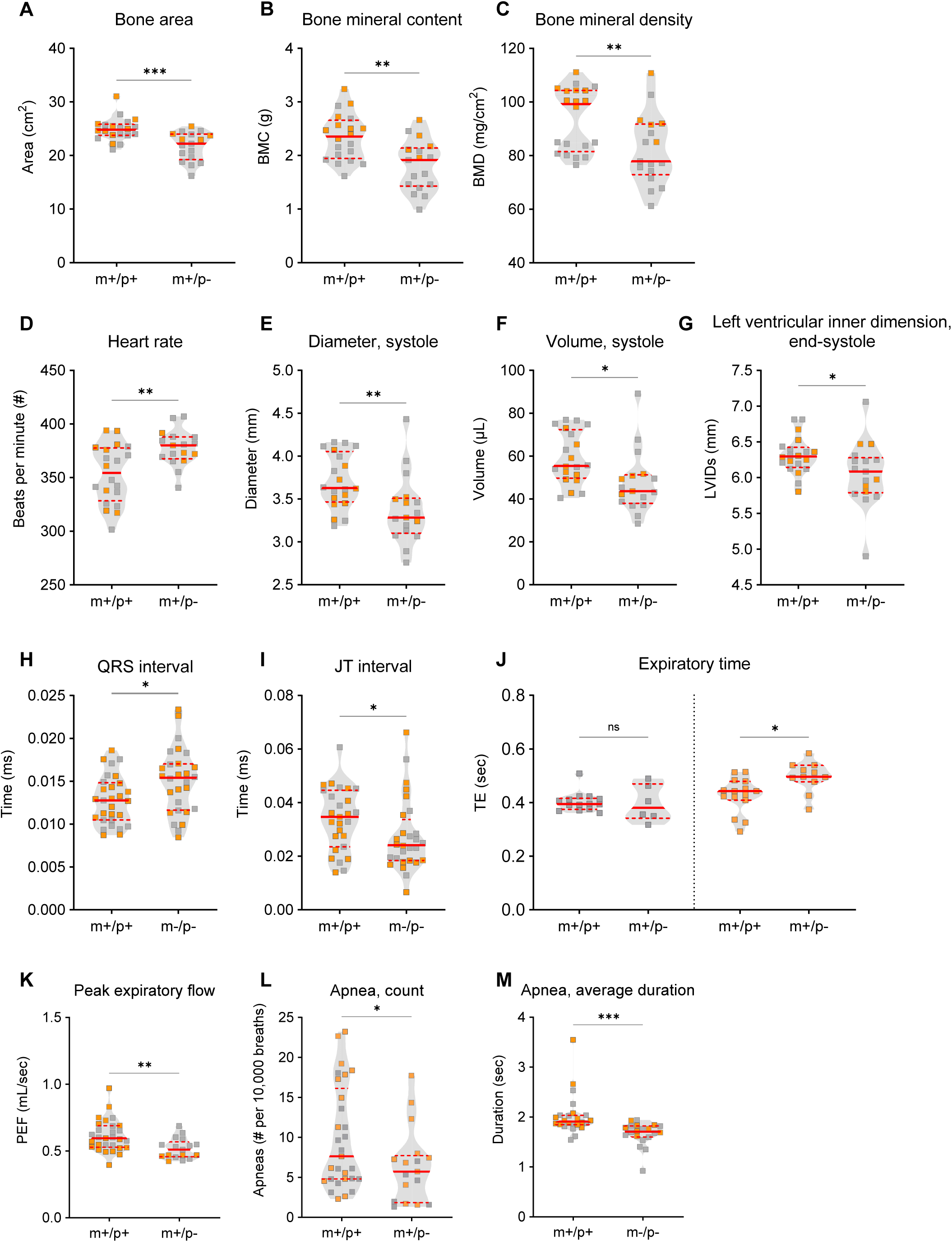
Summary heatmap representation of mutant *Magel2* rat physiology and plethysmography outcomes. **(A-C)** The analysis of bone area, content and density by DEXA scan showed that *Magel2^m+/p-^* (m+/p-) rats compared to their respective wildtype littermates (m+/p+) had a significant decrease in bone area **(A)**, bone mineral content, BMC **(B)**, and bone mineral density, BMD **(C)**. **(D-G)** Cardiac morphology by echocardiogram analysis revealed a significant increase in heart rate **(D)**, decrease in ventricular diameter **(E)** and volume **(F)** during systole in m+/p- rats compared to m+/p+ littermate controls. End systole left ventricular inner dimension during systole (LVIDs) was also reduced in m+/p- rats compared to m+/p+ littermate controls **(G)**. **(H, I)** *Magel2^m-/p-^* (m-/p-) rats compared to their respective wildtype littermates (m+/p+) showed altered cardiac electrical activity measured by electrocardiography; a significant increase in QRS interval **(H)** and decreased JT interval **(I)** was observed in m-/p- rats. **(J-M)** In unrestrained whole-body plethysmography, both m+/p- and m-/p- rats showed alterations in various breathing-related parameters. While expiratory time was increased in female but not male m+/p- rats **(J)**, genotype differences as a group were observed for peak expiratory flow (PEF) **(K)**. Decreased apnea count was observed in m+/p- rats **(L)**; in contrast, the average duration of apneas was decreased in m-/p- rats **(M)**. Violin plots were used to indicate data density and distribution, shown are grey squares representing males and orange squares representing females within each genotype/group category. Solid red line and dashed red lines indicate median and quartiles; respectively. Asterisks indicate statistical significance of mutant rats relative to wildtype littermate controls. *, *p* <0.05; ** *p* < 0.01; *** *p* < 0.01; ns, not significant. Statistical summary of organ systems physiology and plethysmography data are provided in **Supplementary Material, Table ST2**.

### Select cardiac structural and functional dysfunction is observed in *Magel2* mutant rats

Cardiac function was evaluated through echocardiography and electrocardiography tests, which measure the functionality and electrical properties of the heart. To examine the health of cardiovascular systems of *Magel2^m+/p-^* rats, we performed echocardiograms at a one-month time point. A rodent echocardiogram enables ultrasound-based detection of the chambers of a heart, which can then be analyzed for potential abnormalities (Liu and Rigel, 2009). *Magel2^m+/p-^* rats displayed an increased heart rate (**Fig. 6D**), concomitant with systolic deficiencies defined as the phase of heart muscle contraction to pump blood to the body. These impairments included decreased diameter and reduced volume during systole (**Fig. 6E, F**) and decreased diameter of the left ventricle during end-systole (**Fig. 6G**). In contrast, although *Magel2^m-/p-^* rats were normal across structural parameters, several phenotypes by electrocardiography were observed including elevated QRS interval (**Fig. 6H**) which corresponds to the time taken from the initial depolarization of the ventricle to the repolarization for the next contraction, and decreased JT interval (**Fig. 6I**), which correspond to a decrease in time taken to ventricular repolarization.

### Paternal and homozygous mutant *Magel2* mutant rats have altered respiration including altered apnea-based measurements

It has been recently reported that up to 71% of patients with SYS experience some form of respiratory distress (McCarthy et al., 2018). Due to this increased prevalence of respiratory features in SYS, *Magel2* rats were also examined for respiratory performance using unrestrained whole-body plethysmography. *Magel2^m+/p-^* rats exhibited increased expiratory time, impacting peak expiratory flow (**Fig. 6J, K**). Furthermore, both *Magel2^m+/p-^* and *Magel2^m-/p-^*rats exhibited decreased apnea counts and duration (**Fig. 6L, M**). A heatmap (**Fig. 7**) summarizing the degree of statistical significance for organ physiology outcomes identified in each *Magel2* mutant test cohort relative to their respective wildtype littermates is provided, analyzed by multivariate two-way ANOVA for body composition and cardiac measurements (**Fig. 7A**), and for breathing parameters (**Fig. 7B**). Corresponding statistical summary data are provided in **Supplementary Material, Table ST2.**

**Fig. 7.**
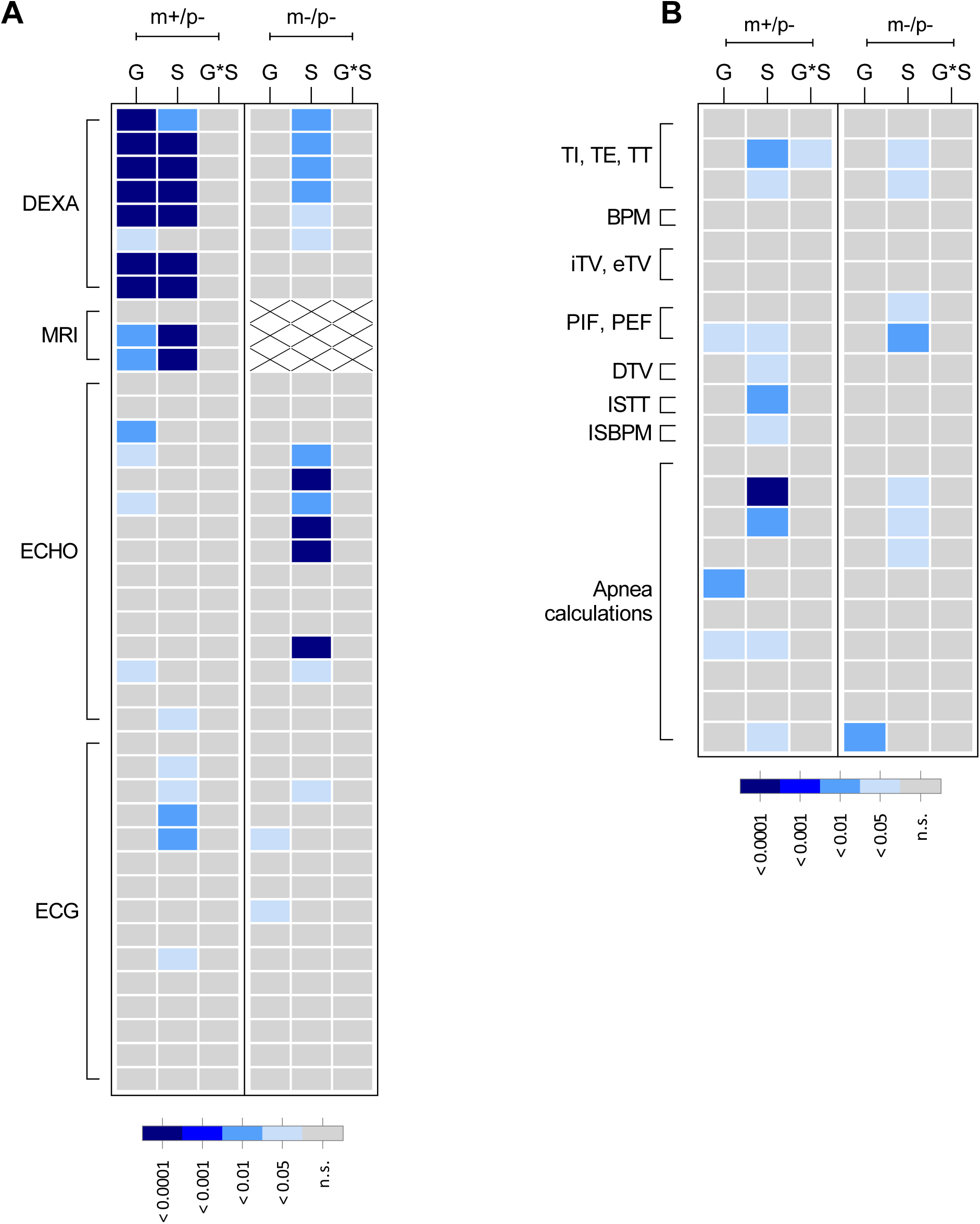
Summary heatmap representation of mutant *Magel2* rat physiology and plethysmography outcomes. Graphical representation showing an overview of the degree of statistical significance for both mutant *Magel2* lines studied relative to their wildtype littermate controls. Each row corresponds to a specific test parameter of the indicated procedure to the left outlined in brackets. Within each mutant line studies (shown along top of graph as m+/p- or m-/p-) each column represents the main effect or interaction for each statistical analysis. As shown in each panel legend, the degree of statistical significance is indicated by relative grey to dark blue color, with each color corresponding to a P value ranging from not significant (ns) to p < 0.0001. For MRI, white boxes with ‘x’ indicate not measured. **(A)** MANOVA with post-hoc analyses were conducted and graphically displayed for dual-energy x-ray absorptiometry (DEXA), quantitative magnetic resonance technology (MRI), echocardiogram (ECHO), electrocardiography (ECG). **(B)** MANOVA with post- hoc analyses were conducted and graphically displayed for multiple parameters collected during unrestrained whole body plethysmography. The specific parameters for each row in both panels **A** and **B** are indicated in **Supplementary Material, Table ST2**; G = genotype, S = sex, G*S = genotype-by-sex.

### Detection of wildtype and novel truncated rat MAGEL2 peptides by mass spectrometry

Detecting endogenous MAGEL2 through antibody-based approaches has been notoriously difficult. Cellular studies have successfully visualized modified MAGEL2 species through genetic approaches such as GFP and Myc labeling (Hao et al., 2013). Recently, a polyclonal antibody was generated against the C-terminal of MAGEL2 and successful in delineating the discrepancy between wildtype and *LacZ* knock in *Magel2* mice, as well as in SYS cell lines (Chen et al., 2020b). However, this antibody would be unable to identify if a truncated MAGEL2 persists in either our novel rat model or in SYS patient cell culture, as the N-terminus is predicted to be absent in truncating mutations.

To determine the presence or absence of wildtype MAGEL2 in our novel rat model we utilized high performance liquid chromatography mass spectrometry. Hypothalamic brain samples from *Magel2^m+/p-^*, *Magel2^m-/p+^*, *Magel2^m-/p-^*, and wildtype littermates were run on an SDS-PAGE gel, a region between 93-170kDa was excised (wildtype rat MAGEL2 predicted molecular weight is 132kDa), gel purified, trypsin digested, and analyzed by mass spectrometry in attempt to identify the full MAGEL2, a truncated species, or if no protein was present across all genotypic combinations (**Fig. 8**). The 8bp deletion generated a frameshift event, p.Ser132Glys, which is predicted to prematurely terminate at amino acid 843 possibly yielding a 71.5kDa truncation product (**Fig. 8A**). From a list of predicted trypsin digested peptide sequences was generated from the predicted translation of wildtype *Magel2*, the most strongly detected peptide sequence from all wildtype littermate samples derived from paternal, maternal, and homozygous inheritance mating schemes was NLPASSETFPATSR. This digested peptide is a translation of nucleotides (c.2725_2766, NC_005100.4), which is 1,982bp downstream of the 8bp deletion and 188bp after the mutant premature stop codon, thus only detectable from a wildtype allele. Ion scores for the wildtype peptide from each sample were collected and suggest the wildtype digested peptide was not detectable in *Magel2^m+/p-^* nor *Magel2^m-/p-^* hypothalamic samples but was detected in *Magel2^m-/p+^* samples (**Fig. 8A, B**). This suggests that i) a full MAGEL2 translation product is very unlikely in any of the mutant states (leakiness is not occurring from the imprinted maternal allele) and ii) maternal heterozygous animals have detectable wildtype peptides. However, a truncated protein could not be empirically discounted. To search for the existence of a truncated MAGEL2 product the mutant *Magel2* sequence was used to generate a list of possible trypsin digested peptide products and a region between 53-93kDa on an SDS-PAGE gel was excised. Three mutant peptides aligning between the 8bp deletion and the premature stop codon, were detected in only *Magel2^m+/p-^* and *Magel2^m-/p-^* samples with high ion scores (**Fig. 8C**). The mutant peptides were not detected in any wildtype samples and did not align to the translated wildtype *Magel2* sequence. Samples from *Magel2^m-/p+^* hypothalamus did not yield detectable mutant peptide sequences, complementing our previous cDNA sequencing data (**Fig. 1**) and further demonstrating that rat *Magel2* is paternally expressed. We report, to our knowledge, of the first protein-based detection of endogenous and unmodified wildtype MAGEL2 along with evidence suggesting the presence of a truncated MAGEL2 species resulting from the *Magel2* mutation in paternal and homozygous mutant rat hypothalami.

**Fig. 8.**
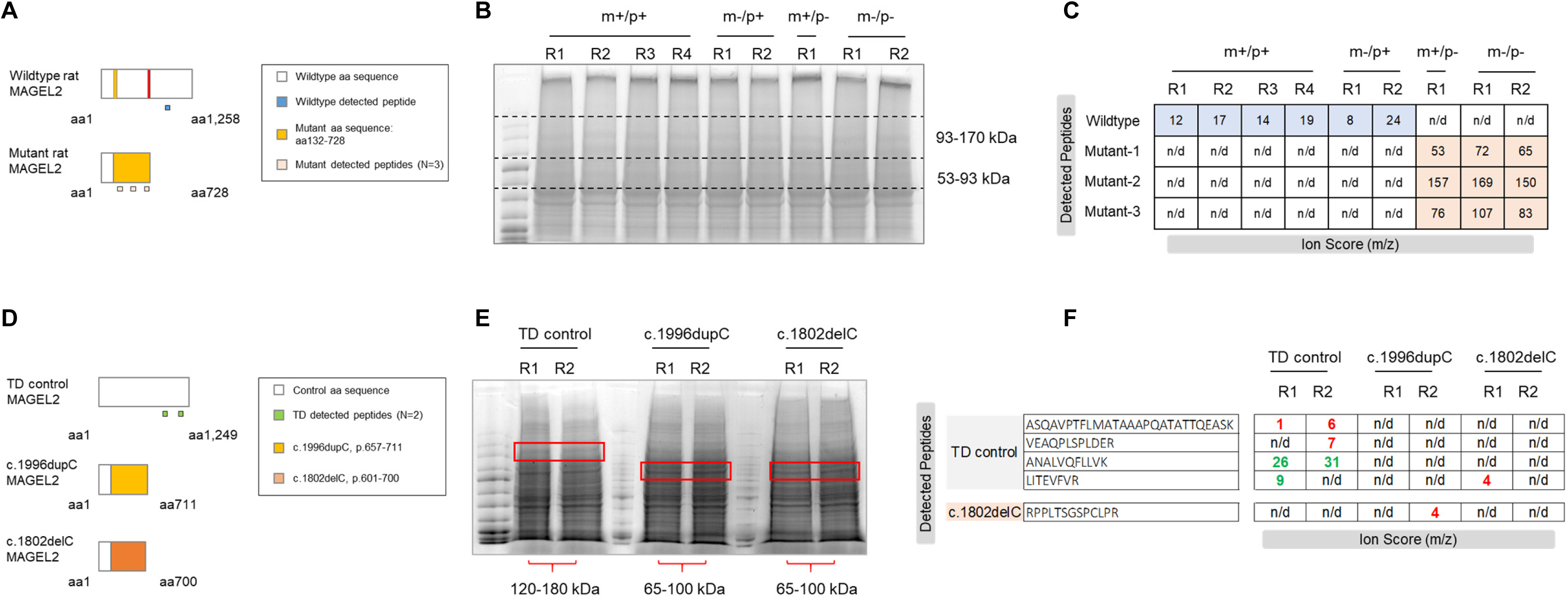
Detection of MAGEL2 peptides in *Magel2* mutant rats and SYS hiPSC lines. **(A)** Schematic diagram showing the expected full-length wildtype MAGEL2 protein compared to the truncated mutant MAGEL2 protein. Within each schematic, the white box indicates wildtype amino acid (aa) sequence; the gold box indicates the mutant aa sequence. The anticipated detection of wildtype (blue boxes) and mutant (light red boxes) peptides are shown below each schematic protein diagram. **(B)** Representative image of SDS-PAGE gel containing wildtype (m+/p+), maternally-inherited *Magel2* heterozygous (m-/p+), paternally-inherited *Magel2* heterozygous (m+/p-), and homozygous (m-/p-) hypothalamus samples with excised regions from 93- 170kDa and 53-93kDa. **(C)** Ion scores (m/z) of detected wildtype and mutant peptides within each allelic combination. R corresponds to the each biological replicate. Rat wildtype MAGEL2 amino acid sequence corresponding to N=1 wildtype peptide was detected in m+/p+ and m-/p+, and not detected in m+/p- or m-/p- rat hypothalami. All three predicted mutant peptides were only detected within the novel region (p.132-728) in m+/p- and m-/p- rat hypothalami. n/d, not detected. **(D)** Schematic diagram showing the expected full-length MAGEL2 protein in typical developing (TD) control hiPSC lines in comparison to the predicted truncated mutant MAGEL2 protein in either c.1996dupC or c.1802delC hiPSC lines derived from SYS individuals. Within each schematic, the white box indicates wildtype amino acid (aa) sequence; the gold box indicates the mutant aa sequence for c.1996dupC, and orange indicates the mutant aa sequence for to c.1802delC. The anticipated detection of TD-detected peptides (green boxes) are shown below each schematic protein diagram. **(E)** Representative image of SDS-PAGE gel containing TD control, c.1996dupC and c.1802delC samples; R indicates biological replicate number. Red boxes outline the excised regions from 120-180kDa (TD control hiPSC) and 65-100kDa (SYS mutant hiPSC). **(F)** Ion scores of detected TD control and mutant peptides within each sample replicate, not detected (n/d), green (strong/confident peptide detection), red (poor/low confident peptide detection). Both TD control replicates had detectable MAGEL2 peptide sequences while neither c.1996dupC nor c.1802delC replicates displayed detectable control or novel mutant sequences.

### Rat and SYS predicted truncated MAGEL2 are similar in size, but human peptides are not detectable in SYS hiPSC protein lysates

Mass spectrometry results were particularly interesting in relation to SYS, whose patients are predicted to have truncating mutations in *MAGEL2* (Schaaf et al., 2013). We asked if the truncated protein generated in our novel rat model was comparable to those predicted in SYS. An extensive report of SYS patients revealed a “mutational hotspot” from c.1990-1996 in *MAGEL2* where the majority of molecularly confirmed cases of SYS reside (McCarthy et al., 2018). 61 of the 78 reported cases were for the mutation c.1996dupC. This prolific mutation is predicted to generate a predicted truncated protein of 72kDa terminating before the USP7- binding (U7B) and MAGE homology (MHD) domains. Upon comparison, the 71.5kDa mutant rat MAGEL2 truncated protein is nearly identical in size and terminates before its predicted U7B and MHD regions.

To determine if predicted truncating mutations in human *MAGEL2* result in a truncated protein, we examined hiPSC SYS cell lines and typical developing (TD) controls with targeted mass spectrometry. A similar approach to distinguishing wildtype and mutant rat MAGEL2 peptides was utilized for the human MAGEL2 sequence (**Fig. 8D**). The SYS hiPSC samples were comprised of either the c.1996dupC or the c.1802delC frameshift mutations. TD and SYS protein lysates were SDS-PAGE separated followed by gel excision of regions predicted for either wildtype or mutant MAGEL2 protein sizes (**Fig. 8D, E**). The c.1996dupC is predicted to result in a mutant peptide sequence beginning at amino acid 666 through its termination at 711, whereas the c.1802delC frameshift occurs at amino acid 601 and terminates at 700 (**Fig. 8D, E**). Mass spectrometry results indicate that wildtype human MAGEL2 peptides are detectable in the TD controls samples while neither wildtype peptides prior to the frameshift mutations nor novel mutant sequences were detected in either of the SYS hiPSC lines (**Fig. 8F**).

## DISCUSSION

Utilizing the TALEN approach (Meek et al., 2017), an eight-base pair deletion was generated in the single coding exon of the rat *Magel2*, resulting in a frameshift event followed by a downstream premature termination codon. This novel mutation was predicted to generate a truncated MAGEL2, separating itself from murine models harboring large gene deletions (Kozlov et al., 2007; Schaller et al., 2010). By generating a truncating *Magel2* mutation in a model with unique experimental advantages such as the laboratory rat, our study has revealed several common and distinct murine alterations in neurobehavioral outcomes and functional changes in organ physiology. F1 hybrid and Sanger sequencing studies confirmed paternal transmission and expression of rat *Magel2,* providing additional confidence in a conserved regulatory mechanism of expression across rat, human and mouse loci (Boccaccio et al., 1999) in typical developing animals. While the Sanger sequencing of the endpoint PCR product across the 8 bp deletion in the *Magel2* rat model do not reveal maternal ‘leaky’ expression, full gene deletions in mice have been reported to release the maternal copy of *Magel2* and other imprinted loci to a limited extent (Matarazzo and Muscatelli, 2013; Rieusset et al., 2013). Future studies examining the methylation of the maternal allele may further elucidate genomic imprinting in the rat, and if truncating mutations may result in low-level maternal expression undetectable in this current study.

At the transcript and protein levels, our analyses of hypothalamic tissue and hiPCs were instructive. Unlike full *Magel2* deletions models which lack endogenous *Magel2* transcript, normal levels of mRNA abundance were observed across all *Magel2* genotypes regardless of uni- and bi-parental contribution. This observation has been previously reported in other monogenic rat models with normal levels of transcript in the presence of altered protein expression (Veeraragavan et al., 2016). To our surprise, by mass spectrometry both wildtype and mutant-derived peptides were detected from rat hypothalamic protein samples but not hiPSCs from SYS and typical developing controls. Several possibilities may explain these results compared to the observations in rat: i) mutant MAGEL2 levels may be extremely low making it difficult to detect via mass spectrometry, ii) the samples were not neuronal cultures (only hiPSCs) whereas *MAGEL2* is neuronally enriched, and iii) the mutant protein may naturally undergo degradation and is not present actively in these samples. Future studies examining the extent of MAGEL2 presence in SYS samples may provide deeper insight into the cellular consequences of this truncated protein. While truncated MAGEL2 protein has still only been predicted to exist in patients with SYS, it is possible that this novel rat model using hypothalamic brain samples may be used as a model to study truncated MAGEL2 as predicted in SYS. These results may further support the hypothesis of a neomorphic truncated mutant MAGEL2, which may explain the observed clinical severity between SYS and full *MAGEL2* PWS deletion.

A primary goal of the comprehensive phenotypic analyses was to evaluate for phenotypic consequences in rats with a truncating *Magel2*. Most literature on *MAGEL2*-related disorders is derived from young children (Patak et al., 2019) with limited studies of the adult phenotype in SYS (Marbach et al., 2020). Our neurobehavioral analysis focused on a neurological comparable rodent age of three to six weeks of age (Semple et al., 2013). Deploying a thorough and comprehensive behavioral battery across multiple domains, we found that *Magel2^m+/p-^* and *Magel2^m-/p-^* rats displayed several alterations from wildtype littermates. *Magel2^m+/p-^* rats exhibited reductions in marble burying during a marble interaction test, suggesting anxiety-like features reflective of a neophobic, avoidant response (Thomas et al., 2009). These data are further strengthened by findings across *Magel2^m+/p-^* and *Magel2^m-/p-^* rats. While *Magel2^m+/p-^* rats appear to have a non-significant reduction in anxiety-like behavior measured in the elevated circle maze, *Magel2^m-/p-^* clear displayed increased time in the open regions of an elevated circle maze. In addition, the play-specific sociability phenotypes observed in the *Magel2* rats points to disruption of social processes systems that may govern direct interaction with respect to time and frequency of social engagement. Social processes in individuals with MAGEL2-related disorders are disrupted as ASD and social deficits are frequently reported (Patak et al., 2019); reduced sociability has also been reported in PWS when compared to individuals with Down syndrome (Rosner et al., 2004). These results appear to indicate that *MAGEL2* may be responsible for select systems within the social processes domain. Finally, while prepulse inhibition was normal across all genotypes, *Magel2^m+/p-^* show a statistically significant reduction in the startle response. *Magel2^m-/p-^* rats in contrast did not show statistically significant differences in either startle reactivity or prepulse inhibition, although it is worthwhile to note that a trend in reduced startle reactivity was observed. Based on the startle circuit, the concordance MAGEL2 disruption may be specifically tied to long-range projections that connect across brain regions responsible for PPI and startle reactivity.

Although different individual sets of phenotypes were uncovered between *Magel2^m+/p-^* and *Magel2^m-/p-^* rats, examination from the perspective of broader phenotypic changes (**Fig. 4**, **Fig. 7**) provides additional insight into how *Magel2* dysfunction in the rat may modulate specific domains especially within the framework of Research Domain Criteria (RDoC) (Insel et al., 2010). Within this perspective of categorical alterations comprised of units of analyses, the results from the *Magel2* mutant rat model implicates altered negative valence behaviors (anxiety-like and perseverative behaviors represented by measures derived from marble burying and elevated circle maze) and altered social processes (represented by measures derived from three-chamber and direct social interactions tests). This interpretation would also be consistent with findings from paternal mutant mice reported to display reduced levels of anxiety-like behaviors in an elevated plus maze and alterations in sociability (Fountain et al., 2017; Meziane et al., 2015). Endophenotypic change within the negative valence domain such as changes in elevated circle, and marble bury are generalized as modulating compulsive/anxiety-like behaviors (Kremer et al., 2021). That people with PWS (Goldstone, 2004; Skokauskas et al., 2012) and SYS (Marbach et al., 2020; McCarthy et al., 2018) display changes in negative valence as rates of anxiety and compulsive behaviors are increased compared to people with non-PWS intellectual disabilities further strengthens the interpretation of these data from an RDoC-based framework.

The second purpose of this study was to evaluate the organ physiology and potential consequences that may arise from a truncating rat *Magel2* mutation. Many individuals with MAGEL2-related disorders are afflicted with debilitating and disruptive changes outside of the central nervous system. The rat is positioned to be an excellent tool for peripheral physiology studies given their size, large tissue area and are a preferred system for therapeutic validation (Wöhr and Scattoni, 2013). From our studies of body composition, cardiac function and structure and respiration, we identified several co-occurring features due to the loss of MAGEL2 in the rat. First, disruption of *Magel2* in rats leads to alterations in body composition highlighted by reduced body weight, whereas gross obesity was never observed. However, findings from *Magel2* mouse models have identified similar metabolic phenotypes in fat mass and body weight, but also have induced obesity, although via a high fat diet (Arble et al., 2016; Bischof et al., 2007). These novel rats were maintained on a standard rodent chow, so the possibility remains that a high fat diet may yield similar results to mouse models. It is worth noting that people with truncating *MAGEL2* variants do not display the hallmark PWS hyperphagia-induced obesity (Chen et al., 2020a; McCarthy et al., 2018). However, a majority of cases within small cohort of adult patients (n=5 of 7) were clinically obese with food-seeking behaviors as reported by caregivers (Marbach et al., 2020). Taken together, the murine model findings strengthen the notion that disruption in *Magel2* across species results in alterations in body composition. Second, through echocardiography, enabling examination of the left ventricle of the heart (Liu and Rigel, 2009), *Magel2^m+/p-^* animals exhibited elevated heart rates and systolic dysfunction. These bidirectional cardiac changes may be a compensatory mechanism to maintain normal blood flow as the ejection fraction in these animals was normal. A possible explanation for these dual observations may be a compensation mechanism, to maintain regular blood output with a smaller left ventricular diameter and mass, the heart may compensate by increasing the frequency of contractions. Further, *Magel2^m-/p-^* rats when examined by electrocardiography, enabling the detection of electrical activity of the heart, displayed an elongated QRS interval, suggesting the time to depolarize the heart was altered. Increase in the QRS interval have been attributed in bradycardia (reduced heart rate)(Da Costa et al., 2002). Additionally, the JT interval, a normative measurement of ventricular repolarization, was reduced in *Magel2^m-/p-^* rats. Together, the electrical heart activity appears modulated without affecting the livelihood of the animals. Relevant to SYS, fatal cases of congenital heart disease have been reported in *MAGEL2* patients (Chen et al., 2020a), making these cardiac findings especially relevant. Third, the findings that *Magel2* mutant rats experience an altered apnea metrics compared to wild-type littermates is interesting when compared to elevated rates of obstructive sleep apnea in SYS (Powell et al., 2020). Although respiration was measured in the unrestrained, whole animal during periods of rest to ensure the breathing measurements were not perturbed by movement, future studies investigating the convergence of sleep behaviors and respiratory performance in these rats would be instructive relative to breathing abnormalities as surrogate translational markers for SYS and PWS.

In conclusion, the results reported here demonstrate that the laboratory rat is a viable model for which to study the molecular, behavioral and organ function implications resulting from a truncating mutation in *Magel2*. The conservation of genomic imprinting of the *Magel2* gene across rat and human allows for another preclinical tool in the effort to provide future therapies to a growing patient population. Given the nature of the genetic manipulations in mouse models of total *Magel2*-deficiency, the availability of a truncated rat MAGEL2 protein may provide valuable additional insight into potential molecular discrepancies that exist between the full deletion and truncating genotypes. The novel rat model with a truncated MAGEL2 also presents an under-appreciated challenge for therapeutic strategies in PWS and MAGEL2-related disorders. While preclinical studies in gene-based interventions have demonstrated the promise of targeted approaches in modulating gene levels, especially for imprinting disorders such as Angelman syndrome (Meng et al., 2015; Wolter et al., 2020). However, in the case of truncating mutations in *MAGEL2*, future gene-based therapy efforts will need to take careful considerations when attempting to activate the silent maternal copy along with possibly targeting the existing pathogenic truncated MAGEL2 species through techniques like proteolysis-targeting chimeras (PROTACs)(Schapira et al., 2019). This current rat model may serve as a useful benchmark along with full gene deletion mouse models in identifying the best approaches for wild-type restoration and reduction of the mutant protein. Overall, these data in the rat model and those reported in mice, link the loss of wild-type MAGEL2 with phenotypes consistent with altered negative valence and social processes, and consistent with alterations observed in people with MAGEL2-related disorders.

## ACKNOWLEDGEMENTS

We thank members of the Samaco lab for technical assistance. This work was supported by the Foundation for Prader-Willi Research (RS), Stedman West Endowed Fund in Neurological Research (RS), and the U.S. National Institutes of Health National Institute of Child Health and Development *Eunice Kennedy Shriver* Baylor College of Medicine Intellectual and Development Disabilities Research Center (grant number P50HD103555) and NIH NICHD R01HD083181 (RS). The content is solely the responsibility of the authors and does not necessarily represent the official views of the National Institutes of Health. The BCM Mass Spectrometry Proteomics Core (AM, AJ) is supported by the Dan L. Duncan Comprehensive Cancer Center NIH award (P30 CA125123), CPRIT Core Facility Awards (RP170005 and RP210227), the Intellectual and Developmental Disabilities Research Center Cores support (P50HD103555), and NIH High End Instrument award (S10OD026804, Orbitrap Exploris 480).

## METHODS AND MATERIALS

### Animals

All animals, studies, and animal care protocols conducted in this study were approved by the Baylor College of Medicine Institutional Animal Care and Use Committee. Rats were housed in a temperature-controlled environment on a 12-h light: 12-h dark cycle with *ad libidum* access to food and water. The *Magel2* mutation was created by transcription activator-like effector nuclease (TALEN) technology which generated the following eight base pair deletion, c.735- 742del, resulting in a frameshift event followed by an early termination site, p.Ser132GlysfsTer728. All rats in this study were bred on a Sprague-Dawley background (Crl:CD-IGS background, strain Code 001, Charles River Laboratories, Inc.). Paternally- inherited *Magel2* heterozygous rats (*Magel2^m+/p-^*, also denoted as m+/p-) and wildtype littermates were generated by mating wildtype females to males heterozygous for the *Magel2* mutation. Homozygous animals (*Magel2^m-/p-^*, also denoted as m-/p-) for the *Magel2* mutation were generated from mating heterozygote females with heterozygote males. Maternally- inherited *Magel2* heterozygous rats (*Magel2^m-/p+^*, also denoted as m-/p+) were generated by mating heterozygous females to wildtype males. At PND14, ear punch biopsies were used for rat identification and PCR genotyping with the following primers: forward, 5’- cagatggcccaatcttcaac-3’; reverse, 5’-tcccaggagggtgtgtcata-3’. PCR products were separated by 4% agarose gel electrophoresis, stained and visualized using an automated gel imaging system (Gel Doc XR+, Bio-Rad, USA). The expected size products for the wildtype allele (115 bp) and mutant allele (107 bp) were used for genotype interpretation. At PND21, rats were weaned and randomly assigned to housing configurations comprised of 2-3 rats per cage. Wildtype animals used in the three-chamber approach and direct social interaction assays were bred from a separate set of mating cages, generating conspecific non-littermate partners.

### Quantitative real time reverse transcriptase PCR (QRT-PCR)

Animals were anesthetized via isoflurane induction, and hypothalami were manually dissected from brain samples and flash frozen in liquid nitrogen. RNA was extracted from dissected specimen using TRIzol reagent (ThermoFischer Scientific, USA). 1-2 µg of total RNA was used for cDNA synthesis; mRNA was selectively reversed transcribed by targeting the poly(A) tail region using only Oligo(dT)_20_ primers (SuperScript IV First-Strand Synthesis System, ThermoFischer Scientific, USA). SsoAdvanced Universal SYBR Green Supermix (Bio-Rad, USA) was used for quantitative real time PCR with commercially available rat *Magel2* and *Gapdh* primers (Bio-Rad, USA; PrimePCR^TM^ SYBR Green assay ID qRnoCED0053164 and qRnoCID0057018; respectively). Expression levels from RT-qPCR experiments utilizing *Magel2* mutant samples were processed as above and normalized to wildtype littermate levels. Statistical analysis of QRT-PCR data was performed using an unpaired t test with genotype as the main factor and graphically presented using GraphPad Prism (Version 9.4).

### Endpoint PCR and Sanger sequencing analysis

To determine if *Magel2* in the rat exhibited parent-of-origin effects, wildtype Sprague-Dawley (Crl:CD-IGS) females and Long Evans (Crl:LE) males were bred to produce F1 hybrid offspring. Hypothalami from parental rats and offspring were dissected for RNA extraction and cDNA synthesis as described above. Endpoint PCR amplification was performed using the primers: forward, 5’- tctaggggccccaatggt -3’; reverse, 5’ ctgctgggctatcggtgt -3’. The expected size PCR product of 460 bp was gel extracted and purified for sequencing analysis (QIAquick Gel Extraction Kit, QIAGEN). Sanger sequencing was conducted by the Baylor College of Medicine DNA Sequencing Core using a 3130XL Genetic Analyzer (Applied Biosystems). Alignment of chromatogram traces to analyze nt position 864 as either G or A was used to determine parent-of-origin inheritance and expression in typically developing rats through the paternal lineage in F1 hybrid test progeny. To confirm the paternal pattern of inheritance and expression in the TALEN-induced *Magel2* rat model, test progeny with uniparental heterozygous (either paternal or maternal heterozygous) and biparental homozygous inheritance of the *Magel2* mutation were generated. Hypothalami from test progeny were dissected for RNA extraction and cDNA synthesis as described above. Endpoint PCR amplification was performed using the genotyping primers which span the eight base pair deletion. PCR product was gel extracted and purified for Sanger sequencing. Presence of wildtype sequence in the chromatogram traces through the *Magel2* mutation site was interpreted as wildtype expression, while presence of a gap in the sequence trace when aligned to a reference sequence was interpreted as expression of the mutated allele.

### Overview of behavioral procedures and statistical analyses of behavioral data

Behavioral performance of juvenile animals from three to six weeks of age was measured using several procedural tests similar to previous studies (Veeraragavan et al., 2016) including anxiety-like behavior, indirect social approach, direct social interaction, locomotor activity, sensorimotor gating, perseverative behaviors, aversive fear memory and object recognition memory, and thermal nociception. Studies were conducted between the hours of 12:00-18:00 (ZT5-11) during the light cycle. Prior to commencing the test battery, each subject was handled for two consecutive days by a trained experimenter blinded to genotype. All assays were performed under dim lighting conditions (8-10 lux) and white noise (60-62 dB) unless otherwise stated. Subjects were habituated to the procedure room for 30 min prior to testing. Both sexes were examined in each behavior cohort with male rats tested chronologically before females in each assay. Group sizes were calculated using a power of 0.8 (alpha=0.05) to detect a difference of one standard deviation based on subjects per genotype per sex. For the behavioral evaluation of paternally-inherited *Magel2^m+/p-^* rat studies, N=12-19 rats per genotype per group were tested. For the behavioral evaluation of homozygous *Magel2^m-/p-^* rat studies, N=14-22 rats per genotype per group were tested. All data were analyzed and represented as individual units within the violin plot formatted graphs. The experimental design strategy deployed for our studies adhere to ARRIVE guidelines (Sert et al., 2020), follows reported study recommendations (Landis et al., 2012), and have been used for the analysis of genetic rat models by our group and others (Berg et al., 2020; Hamilton et al., 2014; Ku et al., 2016; Veeraragavan et al., 2016). *Post-hoc* Tukey comparisons were conducted when appropriate.

Statistical analyses of data were performed using SPSS (Version 28) and graphically presented using GraphPad Prism (Version 9.4). As indicated in **Supplementary Material, Table ST1**, elevated circle maze (ECM), three chamber (3CH), block chewing (BC), novel object recognition (NOR) and fear conditioning training and CS freezing were analyzed using a repeated measures (RM) ANOVA with genotype and sex as main effects. All other behavioral tests and select parameters that did not require statistical modeling using an RM-ANOVA were analyzed using a multivariate ANOVA with genotype and sex as main effects. In cases with significant genotype-by-sex interactions were detected, data were separately analyzed by sex; in the absence of significant genotype-by-sex interactions, genotype comparisons across groups were evaluated.

### Elevated circle maze (ECM)

Animals at the age of 24 days of life were placed on an elevated circular platform with two walled regions (39 cm, height; 60 cm, diameter; 6.3 cm, platform width; 15 cm, closed zone wall height; 46 cm, closed zone wall length). Subjects were placed in the open region, allowed to roam freely, and video recorded for ten minutes. Using the open access event logging software BORIS (Behavioral Observation Research Interactive Software, Torino, Italy), a trained observer blinded to genotypes, scored the recordings for time spent in the closed and open regions. The number of transitions between and the time spent within both regions was exported from the BORIS system. An entrance into either region was defined as a rat having at least half of its anterior torso in the given area.

### Three chamber social approach (3CH)

Animals at the age of 25 days of life were tested in a three-chamber apparatus with perforated walls to allow for interaction between subject and partner rats. A center chamber contained removal doors to two flanking side chambers all with equal dimensions (42.5 cm, length; 17.5 cm, width; 23 cm height). The two side chambers contained a perforated plexiglass wall to allow for indirect contact with novel partner or object behind it. Two sessions were conducted for each subject animal; first, a habitation phase of ten minutes allowing acclimation to the apparatus, and a second ten-minute phase during which a novel conspecific sex matched partner was placed randomly into one of the side chambers with a novel object placed in the other. The subject was allowed ten minutes to freely explore the apparatus and indirect social approach was scored using BORIS by a blinded scorer for time spent in each chamber and time spent directly at each perforated partition.

### Direct social interaction (DSI)

At 26 days of life subjects were single housed overnight in clean standard rat housing cages (40.6 cm, length; 20.3 cm, width; 19 cm, height) with only Sanichip bedding, free of enrichment. The following day, the subject’s cage was placed inside a sound attenuating chamber along with a wildtype same sex conspecific partner introduced. For ten minutes the interaction allowing direct contact was video recorded. A highly trained blinded observer scored the ten-minute interaction for juvenile social play behaviors using the BORIS system. Observations scored included the duration and number of events of play behavior (combination of nape/pounce, wrestling, and pinning behaviors), sniff and following of the conspecific, and paw-to-torso contact towards the conspecific. The aggregate sum of duration and events across all categories were calculated and classified as all active direct social behaviors.

### Open field assay (OFA)

At 27 days of life subjects were allowed to explore an open field apparatus (60cm, length; 60cm, width; 30cm, height) for fifteen minutes. Several parameters were measured via a Fusion v5.3 SuperFlex Open Field System (Omnitech Electronics Inc., Columbus, Ohio) which detects movement by breaks in infrared beams such as horizontal and vertical activity, center distance as well as measurements of time spent in the center of the open field. According to the Fusion v5.3 manual, a 20.32cm x 20.32cm region in the middle of the open field is defined as ‘center’, with the area surrounding it as the ‘margin’.

### Acoustic startle response (ASR) and prepulse inhibition (PPI) of the ASR

At 28 days of life subjects were evaluated for sensorimotor gating using the SR-Lab System (San Diego Instruments, San Diego, CA, USA). Each subject was tested individually; at testing, animals were placed in plexiglass restrainers containing a force meter and positioned within the center of a soundproof chamber. An acclimation phase of five minutes with 70dB white noise preceded each test session. Each test session consisted of 48 trials of randomized presentations of max decibel sound of 120 dB to measure the baseline acoustic startle response (ASR), and presentations of three prepulses of 74 dB, 78 dB, and 82 dB preceding the max decibel sound. Percent PPI of the ASR is calculated as 100-[(prepulse+max startle stimulus response/max startle response alone) x 100], with the average of all three PPI levels used to calculate the global average PPI percentage.

### Marble bury (MB)

At 29 days of life, subjects were placed in a rat cage filled with corn cob bedding (approximately 10 cm) containing twenty marbles arranged in a 4 x 5 matrix and allowed to explore for thirty minutes. After the testing session, animals were removed and the number of marbles visible, half-buried, and three-quarters buried were counted. The number of marbles at least half-buried was used for a measurement of novelty-induced repetitive-like burying behavior.

### Novel object recognition (NOR)

At 30 days of life subjects were habituated to an empty plexiglass open field (60cm, length; 60cm, width; 30cm, height) for five minutes and subsequently placed into an identical open field containing two identical unfamiliar objects (Lego: 6.5cm, length; 6.5cm, width; 16.5cm, height) placed catercorner from each other for another five minutes. Subjects were video recorded during the exposure trial to the two objects. The next day, the same subjects were again habituated in an empty open field for five minutes and placed into the test arena with one of the previous objects replaced randomly with a novel object. The trial session was video recorded, and a blinded scorer recorded the time the subject spent at each of the two objects. The novel object recognition index was determined by the time spent at the novel object versus total time between the two objects [(time at novel object/total time spent at both novel and familiar objects) x 100].

### Fear conditioning (FC)

On days 32 and 33 of life, subjects were tested in a classical fear conditioning paradigm. The animals were placed in a metal bar bottomed chamber (26 cm, length; 30 cm, width; 20 cm, height) for 120 seconds followed immediately by a 30 sec period with 85 dB noise, ending with a 2 sec, 1 mA foot shock. The following day the subjects were placed back into the original chamber for 300 sec to assess contextual fear memory. To evaluate cued fear memory, the chamber was cleaned, and black plexiglass was inserted into the chamber along with an artificial vanilla scent to generate a novel environment. Subjects were returned to the modified chambers for 180 sec followed by conditioned stimulus 85 dB for the remaining 180 sec. Freezing behavior, a surrogate readout of memory, was measured for both contextual and cued fear memory. Freezing behavior is expressed as a percentage of time relative to the total test duration in which the subjects are motionless for >1 sec; this is determined using automated detection software enabling video capture of frame-to-frame changes in movement (FreezeFrame version 4.104, Actimetrics, Wilmette, IL, USA).

### Block chewing test (BC)

At 34 days of life subjects were single housed without environmental enrichment for twenty-four hours with access to a single wooden block (Aspen wood; 0.4 cm, length; 0.2 cm, width; 0.2 cm, height). Weight of the wooden block was measured before and after placement within each a subject’s cage. The difference in wooden block weight was taken to correspond as a measure of perseverative-like behavior.

### Hot plate (HP)

At 35 days of life subjects were tested for pain sensitivity to thermal heat. Subjects were placed on a specialized test chamber containing a plate heated to 55^°^C with a plexiglass chamber surrounding the test area. The time to the first instance of shaking or licking of the hindpaws was recorded, and indicated as the measure of latency to first response. Subjects were immediately removed from the test chamber upon observation of the first response and returned to their housing configuration.

### Overview of peripheral organ system assessments and statistical analyses of data

Rats heterozygous for the paternally-inherited *Magel2* mutation (m+/p-) and rats homozygous with biparental-inheritance of the *Magel2* mutation (m-/p-) were evaluated for alterations in body weight and composition, cardiac structure and function, and breathing patterns. For all physiological measures examined in paternally-inherited *Magel2^m+/p-^* rats except breathing parameters, N=9-11 rats per genotype per group were tested; for breathing studies N=6-14 per genotype per group were tested. For all physiological measures examined in homozygous *Magel2^m-/p-^* rats except breathing parameters, N=11-18 rats per genotype per group were tested; for breathing studies N=10-12 per genotype per group were tested. All data were analyzed and represented as individual units within the violin plot formatted graphs. Statistical analyses of data were performed using SPSS (Version 28) and graphically presented using GraphPad Prism (Version 9.4). As indicated in **Supplementary Material, Table ST2**, all data were analyzed using a two-way ANOVA with genotype and sex as main effects. Parameters showing genotype-by-sex interactions were analyzed with genotype differences examined within sex. In the absence of significant genotype-by-sex interactions, data were analyzed with genotype as the main effect. With the exception of body weight, body composition measures were collected from m+/p- rats at one and three months of age, and m-/p- rats at one month of age.

### Dual-energy x-ray absorptiometry (DEXA)

To ascertain the body composition of rats mutant for *Magel2*, weights were first manually recorded and individual rats were anesthetized with 2.5-3.5% isoflurane with 0.5L/min of 100% O_2_ and placed ventral side down inside of a Faxitron® UltraFocus^DXA^ system with a nose cone to maintain isoflurane flow. The limbs and tail were spread away from the torso to capture clean scans with the spine straightened. Parameters including bone mineral content, bone mineral density, fat weight/percentage, lean muscle weight/percentage, and total weight were collected as an output text file for each rat.

### Quantitative magnetic resonance technology (EchoMRI)

Due to the size of rats at three months of age, DEXA using the Faxitron® could not be performed. Instead, body compositions were acquired through an EchoMRI^TM^ Body Composition Analyzer using quantitative magnetic resonance technology. Weights of each rat were manually recorded before being placed into the EchoMRI^TM^ machine. Each subject rat was placed into and gently guided to end of the size 700-gram holder with a Velcro barrier placed to minimize movement. A noninvasive scan was then performed on each rat while awake to obtain measurements of fat and lean muscle masses.

### High frequency ultrasound echocardiogram (ECHO)

On the day prior to testing, each rat was fully anesthetized with isoflurane, placed on a heating pad to maintain body temperature, and the ventral region from its neck to xiphoid process was removed of hair using Nair^TM^ and a piece of gauze. The subsequent day, rats were anesthetized with 2.5% isoflurane mixed with 0.5L/min of 100% O_2_ and placed ventral side-up on an electric heating pad. Body temperature was continuously monitored throughout the procedure with a lubricated rectal probe and maintained between 36.5-37.5^°^C. Pre-warmed echocardiography gel was applied to the hairless region above the heart. High frequency ultrasound echocardiography was performed using a VisualSonics Vevo® 2100 Imaging System with a MS250 transducer attached (FUJIFILM VisualSonics Inc., Toronto, ON) to acquire ultrasound-based images of the left ventricle. Cross-sectional views of each rats’ heart were generated using the parasternal short-axis mode and data acquired by recording from the papillary muscles.

### Surface electrocardiography (ECG)

To measure the electrical properties of the rhythmic firing pattern of the left ventricle, we utilized an Indus Rodent Surgical Monitor^+^. Rats were anesthetized using 2.5% isoflurane with 0.5L/min 100% O_2_ and positioned ventral side down onto the heated surgical platform. Electrode cream was placed on each of the fore and hind paws, which were taped in place over individual leads. Following each cardiac cycle, amplitude and time of the P, Q, R, S and T waveforms, QT interval, ST segment height were recorded and averaged, along with ventricular repolarization parameters like the JT interval were also calculated. Each rat tested had at least 1,000 cardiac cycles measured, of which the above parameters were averaged across. Arrhythmic activity was monitored during each session by the trained experimenter.

### Whole body unrestrained plethysmography

Rats were brought to the test room, their body weights recorded and were then habituated to the testing room for 30 minutes. After the habituation period, each animal was placed within the plethysmography recording chambers and breathing was recorded with synchronized capture of video for 1 hour using data collection software (Ponemah 5.2 with Noldus Media Recorder, DSI). After testing animals were returned to their home cages. Videos were scored by automated algorithm to identify periods of calm state, and breathing signal were scored by automated algorithm to identify breaths. Summaries of the breath parameters for each animal while in a calm state were generated and used for analysis (Ward et al., 2020).

### Human induced pluripotent stem cell (hiPSC) culture and protein lysate collection

All hiPSCs were maintained on GelTrex (Thermo Fischer Scientific) in mTeSR Plus (Stem Cell technologies) or in StemMACS iPS-Brew XF (Milteny Biotec) with medium change every other day and kept in a 37°C incubator with 5% CO2. hiPSCs were regularly split at around 80% confluency or earlier based on morphology with Versene (Gibco) treatment. For total protein extracts cells were harvested, washed with PBS and lysed in RIPA buffer containing Halt Protease-Inhibitor-Cocktail (Thermo Fischer Scientific) for 1 h on ice. Quantity of protein were assessed by Pierce™ BCA Protein Assay Kit (Thermo Fischer Scientific) according to the manufacturer’s instruction.

### Mass spectrometry for MAGEL2 wildtype and mutant truncated species

Mass spectrometry experiments were conducted through the BCM Mass Spectrometry Proteomics Core. For each animal, a 5mm^3^ section of rat hypothalamus was dissected and flash frozen in liquid nitrogen and crushed to powder with a pestle on a liquid nitrogen chilled steel block. Powdered tissues were lysed on ice for 30 minutes with 400mL RIPA buffer (50mM Tris-HCl pH7.6, 150mM NaCl, 1mM EDTA, 0.25% Sodium doxycholate, 1% NP-40, 0.1% SDS), supplemented with protease (GeneDepot, P3100) and phosphatase (homemade: 10mM NaF, 1mM Na_3_VO_4_, 10mM ß-Glycerolphosphate) inhibitors. The lysates were further sonicated (GE505 Ultrasonic Processor) 3 times with 10” ON and 30” OFF intervals on 20% power on ice and cleared by centrifugation at 21,000 rpm for 20min at 4°C (Eppendorf 5424R Centrifuge). 50µg of supernatant was mixed with NuPAGE LDS Sample Buffer (Thermo Scientific, NP0007) and boiled at 90°C for 10 minutes. The proteins were separated on pre-cast NuPAGE Bis-Tris 10% protein gel (Invitrogen, NP0301BOX). Gels were stained with Coomassie (0.025%Brilliant Blue R-250, 40% MeOH, 7% AcOH) and two bands were excised corresponding to 93-170kDa and 53-93kDa. The expected molecular weight (MW) of wildtype rat MAGEL2 is ∼125kDa with predicted localization within the 93-170kDa region, while the predicted ∼71.5kDa MW of a truncated MAGEL2 would localize to the 53-93kDa region of the gel. The bands were then crushed and digested with 22mL of Trypsin solution (20mL 50mM ammonium bicarbonate and 2mL of 100ng/mL Trypsin, GenDepot, T9600) overnight at 37°C with shaking. For mass spectrometry runs, excised peptides were re-suspended in 5% methanol + 0.1% formic acid solution and measured on an Exploris Orbitrap 480 mass spectrometer (Thermo Fisher Scientific, San Jose, CA, USA).

Predicted expression of *Magel2* protein is very low, therefore, the mass spectrometry method included a DDA experiment (Top 20 most intense ions with a dynamic exclusion duration of 15 sec) and a targeted experiment for predicted peptides of MAGEL2. As empirical data on behavior of peptides from wildtype rat MAGEL2 were not available to evaluate potential best responders, eighteen peptides were chosen based on their theoretical suitability for detection. A possible truncated MAGEL2 species was evaluated based on predicted peptides following trypsin digestion, similar to the wildtype approach. Raw data were searched against a rat RefSeq RefProtDB. The Proteome Discoverer software suite (PD version 2.0.0.802; Thermo Fisher Scientific) was used to search the raw files with the Mascot search engine (v2.5.1, Matrix Science), validated peptides with Percolator v2.05, and provide MS1 quantification through Area Detector Module. All peptide spectral matches for MAGEL2 were manually evaluated.

For hiPSCs, 20µg of cell lysate was resolved on pre-cast NuPAGE Bis-Tris 10% protein gel. Two band regions were cut as follows: band 1 for 60-90kDa region and band2 for 120-170kDa region. In-gel digestion and peptide extraction was carried out as described before. The peptides were analyzed on a nano-LC 1000 system (Thermo Fisher Scientific, San Jose, CA, USA) coupled to Orbitrap Fusion mass spectrometer (Thermo Fisher Scientific, San Jose, CA, USA). The nanoLC column and gradient settings was same as described before. The MS instrument was operated in DDA mode with a targeted experiment for predicted MAGEL2 peptides in WT and mutants. Peptides were selected based on their uniqueness in each of the genotype. The targeted experiment was performed in IonTrap (HCD 32%) with a 1.6m/z isolation window in the quadrupole. The MS raw data was searched against human NCBI RefSeq protein database (updated 2020-03-24) in Proteome Discoverer software suite. Precursor mass tolerance set to 20ppm and fragment ion tolerance to 0.5Da. All peptide spectral matches for MAGEL2 were manually evaluated. The MS1 and MS2 peaks were extracted using Skyline software (MacCoss lab, University of Washington).

